## Supplementary File 1 for "*Trackoscope*: A Low-Cost, Open, Autonomous Tracking Microscope for Long-Term Observations of Microscale Organisms"

### Contents


***Trackoscope* Wiring Diagram** **2**

***Trackoscope* Prototype** **3**

***Trackoscope* Movement Analysis Calculation** **4**

**Customized *Trackoscope* Prototype** **5**

***Trackoscope* Part List** **6**

***Trackoscope* Assembly Instructions** **7**

### *Trackoscope* Wiring Diagram

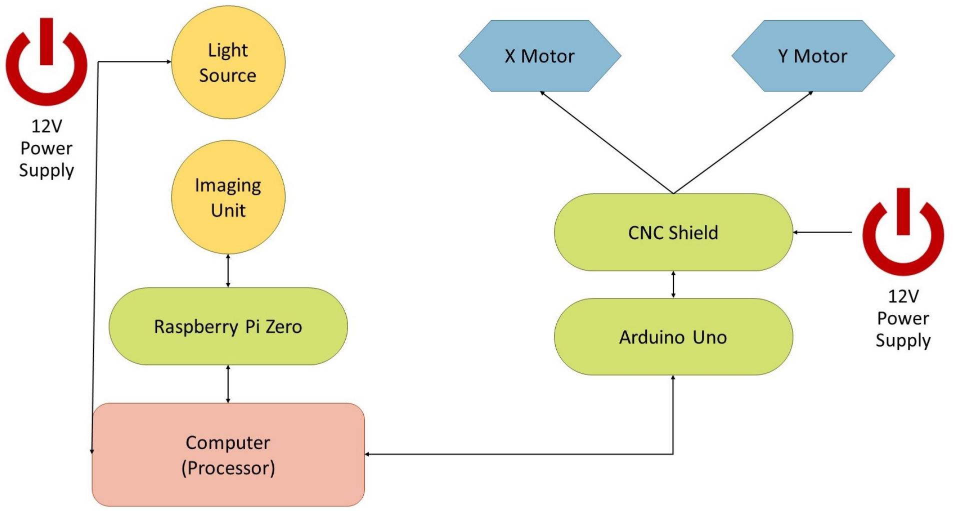


Figure S1: **Circuit Diagram** Wiring diagram for *Trackoscope* detailing the connections between various components.

### *Trackoscope* Prototype

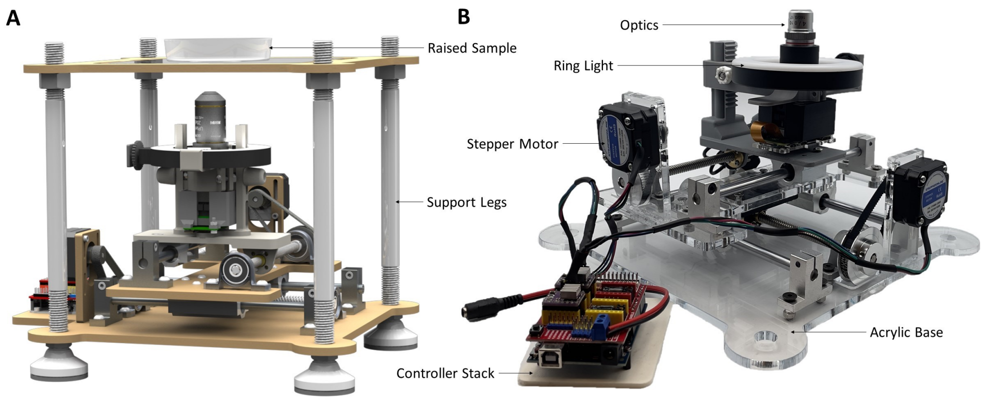


Figure S2: **The *Trackoscope* prototype**. (A) CAD model of *Trackoscope* in the raised sample imaging setup. A white or contrasting covering is typically placed over the sample to create a clean background in the image. (B) *Trackoscope* is designed to be mass-produced and is constructed primarily out of laser-cut parts (acrylic or MDF) and minimal 3D printed components. It also uses standard metric nuts and bolts to join components together.

### *Trackoscope* Movement Analysis Calculation

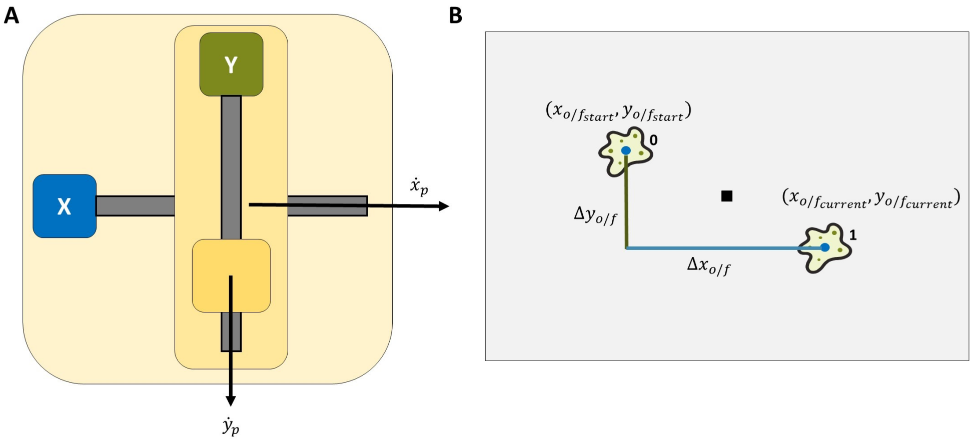


Figure S3:
***Trackoscope* Movement Analysis Calculation**. (A) $(\Delta x_{o/f}, \Delta y_{o/f})$ are calculated by taking the organism's location in the video frame at the start (is not always $(0,0)$) and finding the displacement within the frame, $\Delta x_{o/f}(\mu m) = (\Delta x_{o/f_{previous}} + (x_{o/f_{current}} - x_{o/f_{start}})) * C_{pixels\,to\,\mu m}$, with $C_{pixels\,to\,\mu m}$ depending on the magnification. (B) $(\Delta x_p, \Delta y_p)$ are calculated by adding up all platform displacements throughout the track; for instance, a single data point would be calculated with $\Delta x_p(\mu m) = \Delta x_{previous} + (\dot{x}_p * 50ms)$ where $\dot{x}$ is the velocity of the axis on the platform.

### Customized *Trackoscope* Prototype

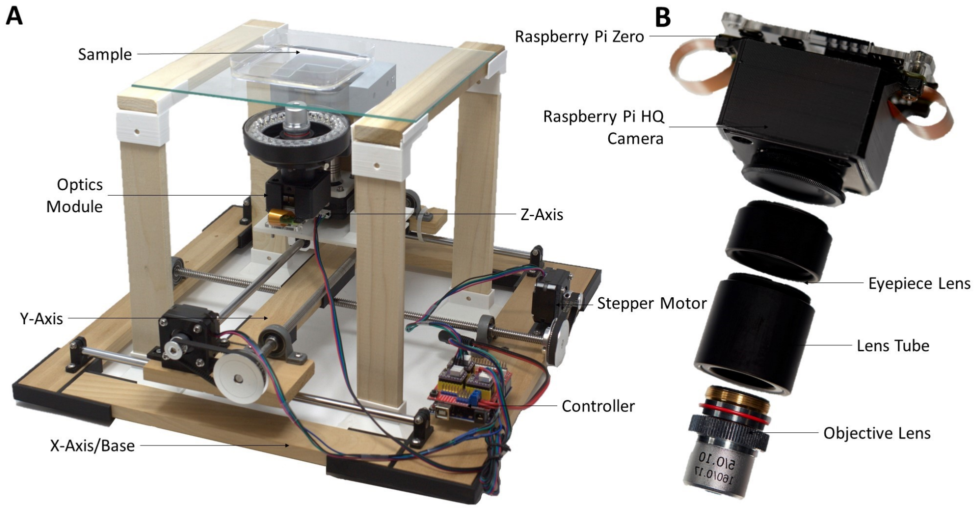


Figure S4:
**Customized *Trackoscope* prototype**. (A) The single-build customized version of *Trackoscope* featuring a motorized Z-axis and a raised sample that is observed from below. This prototype has a tracking area of 625cm$^2$ (size of an A4 sheet) and is built using limited tools (hand saw, 3D-printer, and screw-drivers). (B) The physical custom digital microscope system used in both prototypes.

### *Trackoscope* Part List

| Part | Cost | |
|---|---|---|
| Arduino Uno + CNC Shield (i) | $28.88 | |
| NEMA 17 Stepper Motors (ii) | $25.08 | |
| Threaded Rods Set (iii) | $56.58 | |
| Belt + Pulley (iv) | $21.98 | |
| Legs (v) | $9.99 | |
| Leg Screws (vi) | $3.20 | **Total Cost:** |
| Coupling Nuts (vii) | $6.35 | **$423.54** |
| 3D Printed Parts (viii) | $4 | |
| MDF (ix) | $5 | *Actuator Cost:* |
| Objective Lens (x) | $14.99 | *$167.05* |
| Achromatic Lens (xi) | $110.36 | |
| Ring Light (xii) | $11.99 | *Optics Cost:* |
| Raspberry Pi High Quality Camera (xiii) | $50 | *$256.49* |
| Lens Tube (xiv) | $29.03 | |
| Raspberry Pi Zero (xv) | $15 | |
| Cage Plate (xvi) | $21.11 | |
| Fasteners (xvii) | $10 | |

Table S1: The table lists the cost of *Trackoscope*. (i) a Shield Expansion Board V3.0 + R3 Board + A4988 Stepper Motor Driver + Heatsink, from kumantech.com, P/N K75; (ii) Nema 17 Bipolar 0.9deg 11Ncm (15.6oz.in) 1.2A 3.6V 42x42x21mm 4 Wires, from omc-stepperonline.com, P/N 17HM08-1204S; (iii) Mergorun 200mm Horizontal Optical Axis and 8mm Lead Screw Dual Rail Shaft Support Pillow Block Bearings and Flexible Shaft Coupling, from amazon.com, ASIN B06XPCY1LS; (iv) Houkr PGT2 Aluminum Timing Belt Idler Pulley Bearing 20and60 Teeth Width 8mm Born Synchronous Wheel, with a Perimeter 200mm Width 6mm Belt and a M4 Allen Wrench, from amazon.com, ASIN B081PXKKS4; (v) Antrader Pcs M10 Thread Adjustable Foot Cups Reinforced Nylon Base 48mm Diameter Articulated Feet Furniture Leg 80mm Leveling Foot, from amazon.com, P/N AZ18082701; (vi) 100mm Medium-Strength Metric Class 8.8 Steel Hex Head Screws, from mcmaster.com, P/N : 91280A198; (vii) Hitiland 5pcs Long Rod Nut Carbon Steel Hex Coupling Nuts Hexagonal Sleeve Nut Standoff M10 Threaded Fasteners, from amazon.com, P/N B07QG2YSXT; (viii) 200 grams of PLA filament; (ix) 1/4 in. x 2 ft. x 4 ft. Medium Density Fiberboard, from homedepot.com, P/N 1508104; (x) 4X Achromatic Microscope Objective, from amscope.com, P/N A4X-YX-V460; (xi) AC254-100-C - f = 100.0 mm, Ø1" Achromatic Doublet, ARC: 1050 - 1700 nm, from thorlabs.com, P/N AC254-100-C; (xii) AIXPI 4" Ring Light for Laptop with Stand and Clip, from amazon.com, ASIN B08XQCYPJ3; (xiii) Raspberry Pi High Quality HQ Camera - 12MP, from adafruit.com, 4561; (xiv) SM1 Lens Tube, 0.50" Thread Depth and SM1 Lens Tube, 1.00" Thread Depth, from thorlabs.com, P/N SM1L05 and SM1L10; (xv) Raspberry Pi Zero 2 W, from adafruit.com, P/N 5291, (xvi) 30 mm Cage Plate with Ø1" Double Bore, from thorlabs.com, P/N CP35, (xvii) M3/M4/M6 screws and nuts, from mcmaster.com, P/N 90258A187/90258A221/90258A253/90592A085/90592A090/90592A095 *All links and prices accessed January, 2023.

### Assembly Instructions

#### *Trackoscope* Assembly Instructions

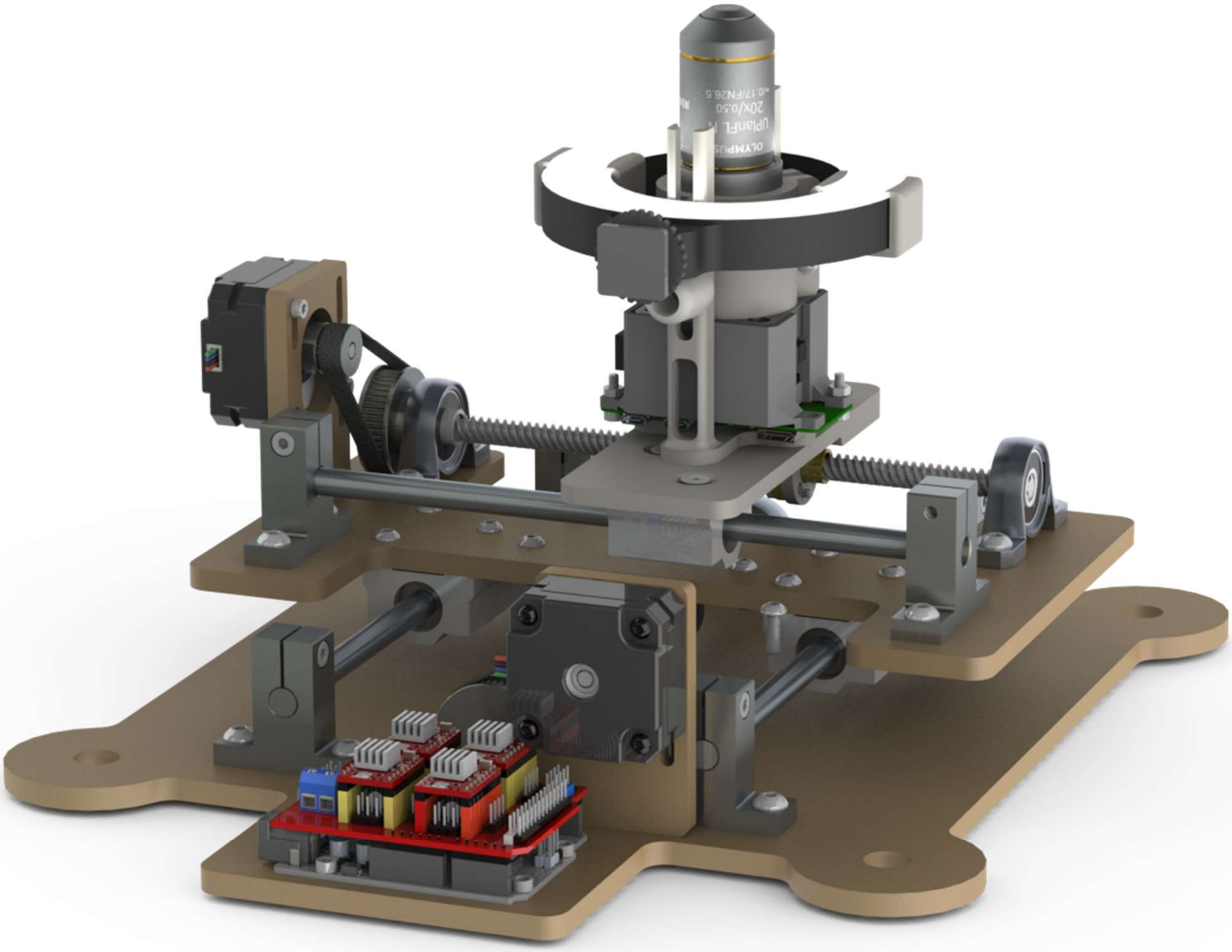

### A. *Optics Module*

#### Parts

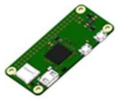 - 1x Raspberry Pi Zero

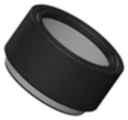 - 1x SM1L05 Lens Tube

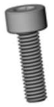 - 2x M3-10 Socket Head Bolt

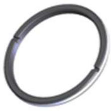 - 1x SM1RR Ring

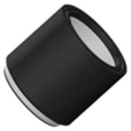 - 1x SM1L10 Lens Tube

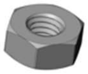 - 2x M3 Nut

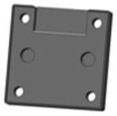 - 1x 3D Printed Top Case

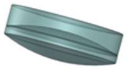 - 1x Achromatic Doublet Lens

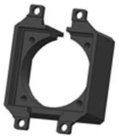 - 1x 3D Printed Bottom Case

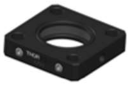 - 1x Cage Plate

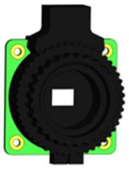 - 1x Raspberry Pi HQ Camera

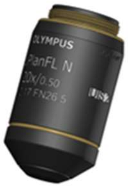 - 1x Objective Lens

| | | |
|---|---|---|
| 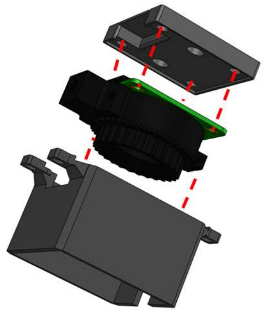 | 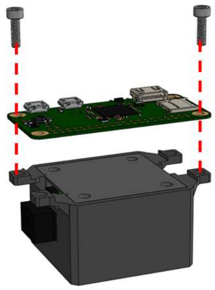 | 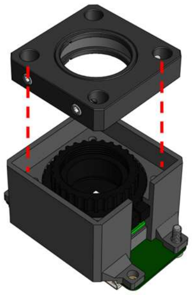 |
| 1. Press the Raspberry Pi HQ Camera inside the two 3D printed pieces | 2. Attach the Raspberry Pi Zero with two M3 bolts. Attach nuts after building | 3. Press the cage plate into the sensor side of the camera |
| 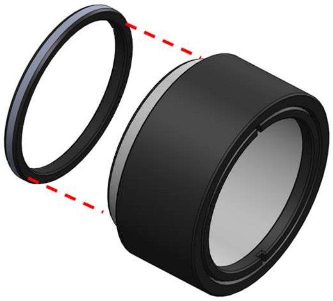 | 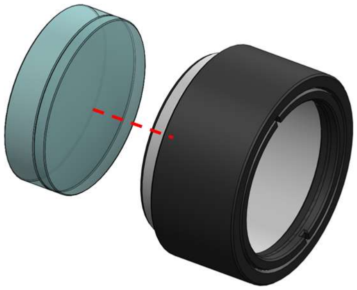 | 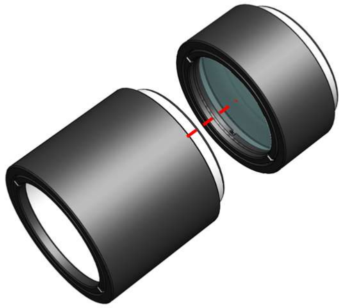 |
| 4. Screw the retaining ring midway into 0.5" lens tube. | 5. Place the lens into lens tube, convex face facing down. | 6. Screw the 1" lens tube to the 0.5" lens tube. Be sure to screw onto the side where the convex face of the lens is. |

| | | |
|---|---|---|
| 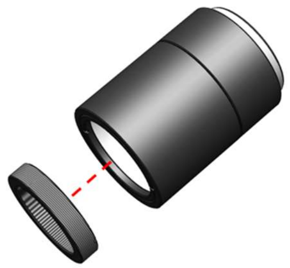 | 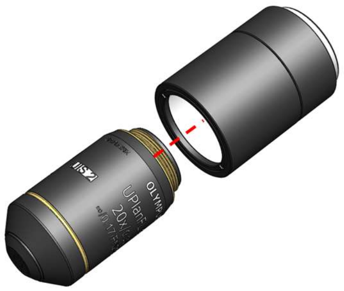 | 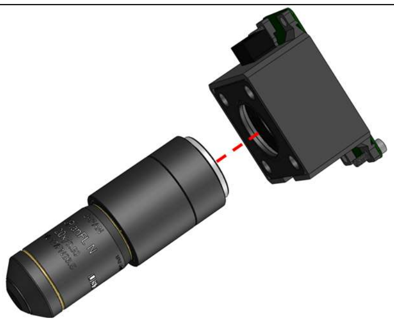 |
| 7. Screw a retaining ring into the lens tube set | 8. Screw an objective lens to the lens tube set | 9. Screw the lens system to the cage plate |

#### *B. Z-Axis Module*

##### Parts

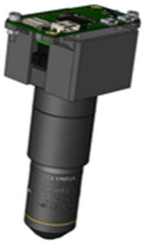

- 1x Optics Module

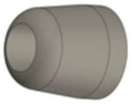

- 2x 3D Printed Knobs

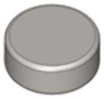

- 10x 6mm Neodymium Magnets

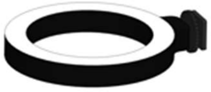

- 1x Ring Light

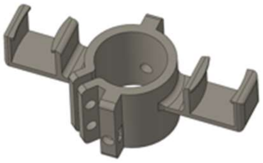

- 1x 3D Printed Combined Hold

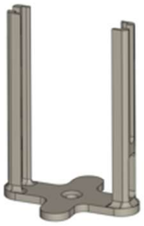

- 1x 3D Printed Stand

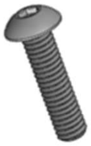

- 4x M4-14 Flat Head Bolt

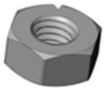

- 4x M4 Nut

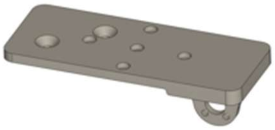

- 1x 3D Printed Base

| | | |
|---|---|---|
| 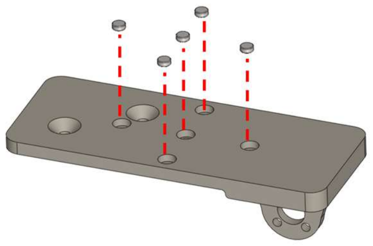 | 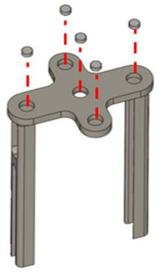 | 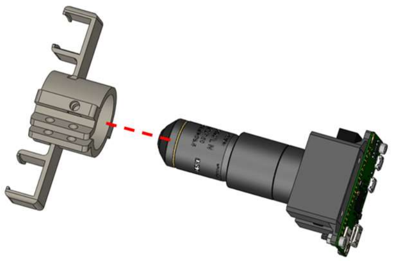 |
| 1. Glue 5 magnets into the base portion (ensure magnet polarity is the same) | 2. Glue 5 magnets into the stand portion (ensure magnet polarity is the same and opposite of base magnets) | 3. Slide the optics unit into the combined hold |
| 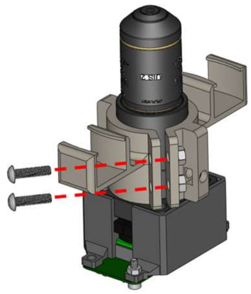 | 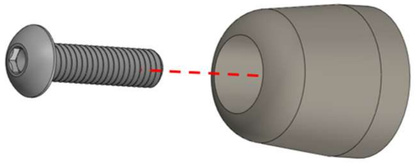 | 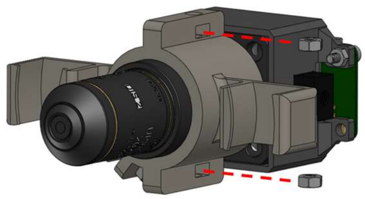 |
| 4. Clamp the optics with 2 M4 bolts and nuts | 5. Glue an M4 bolt into a knob. (repeat twice for the second knob) | 6. Slide 2 M4 nuts into the gaps in the combined hold piece |
| 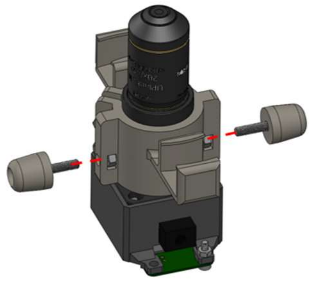 | 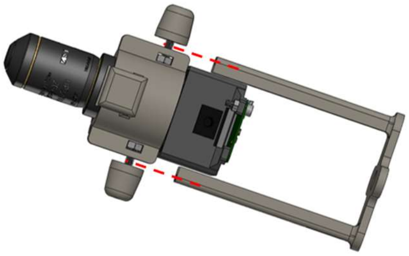 | 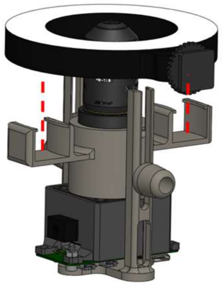 |
| 7. Screw the knobs in slightly till they grasp onto the nut | 8. Slide the combined module into the stand and screw in the knobs till the system is secure | 9. Snap the ring light into the module |

#### *C. Y-Axis Module*

Parts

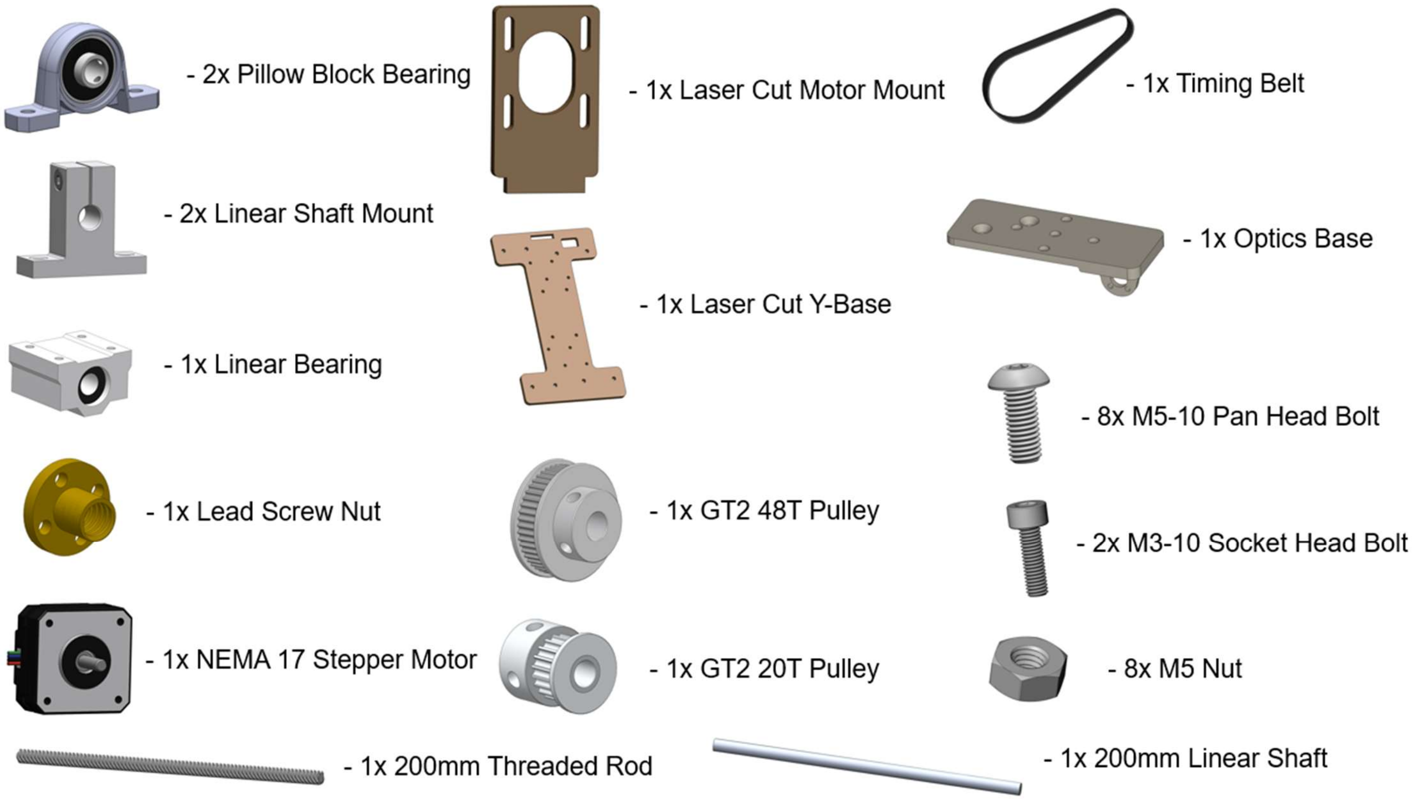


| | | |
|---|---|---|
| 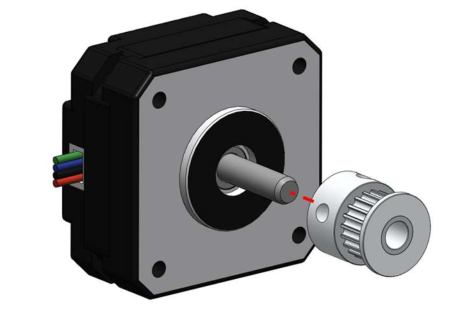 | 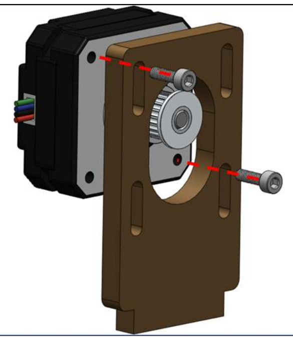 | 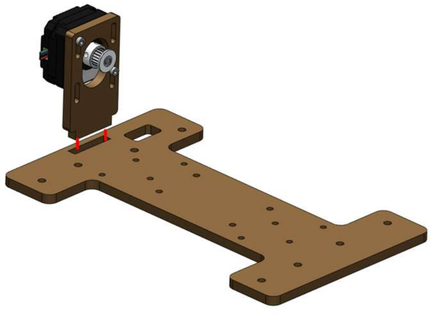 |
| 1. Slide the 20T pulley onto stepper motor shaft. Tighten the two set screws on the pulley. | 2. Screw the stepper motor onto the laser cut motor mount. | 3. Glue the motor mount into the laser cut y-axis base. Use super glue or epoxy. Let fully cure. |
| 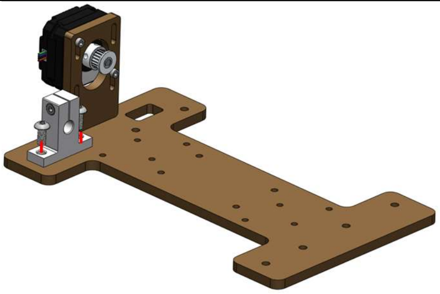 | 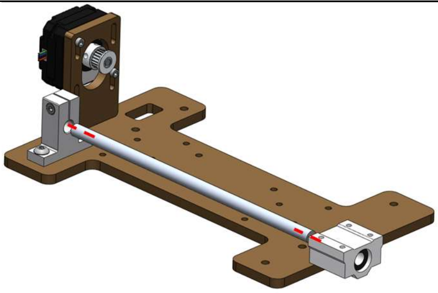 | 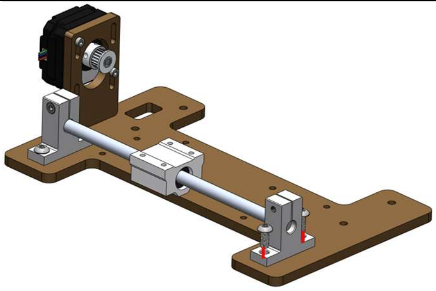 |
| 4. Screw a linear shaft mount onto the base with M5 bolts. | 5. Slide the linear shaft into the mount and slide a bearing onto the shaft. | 6. Screw a linear shaft mount onto the base with M5 bolts. |

| | | |
|---|---|---|
| 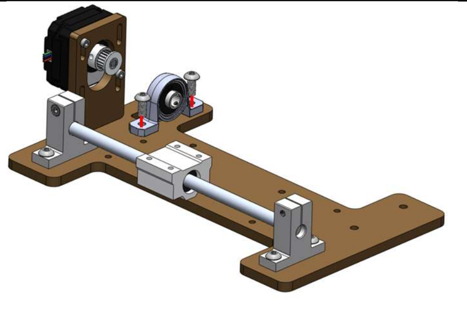 | 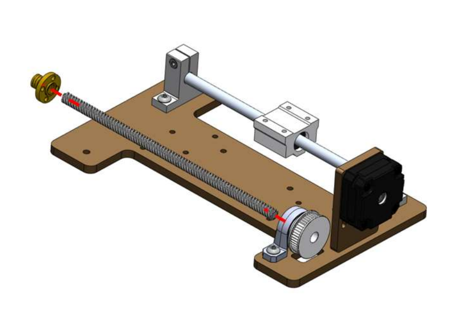 | 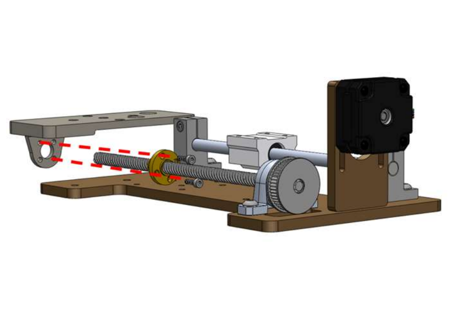 |
| 7. Screw a pillow block bearing onto the base with M5 bolts. | 8. Place the belt on the 42T pulley and slide the threaded rod through the pillow block bearing and the pulley. Spin the lead nut onto the rod. | 9. Screw the optics base to the lead screw nut using two M3 nuts and bolts. |
| 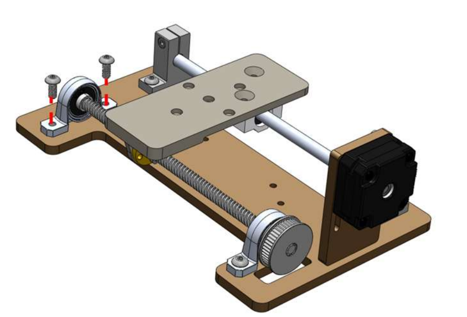 | 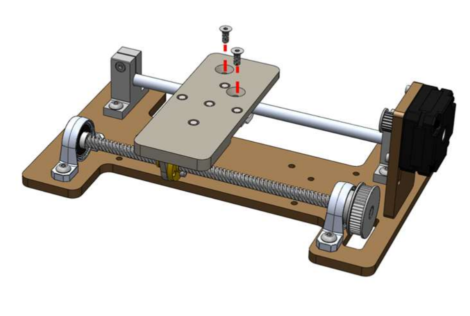 | 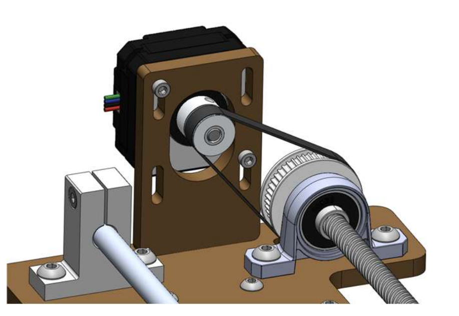 |
| 10. Attach the pillow block bearing onto the base with M5 nuts and bolts. | 11. Attach the optics base to the linear bearing with two M4 nuts. | 12. Adjust the motor mount screws to maintain slight tension in the belt. |

#### *D. X-Axis Module*

##### Parts

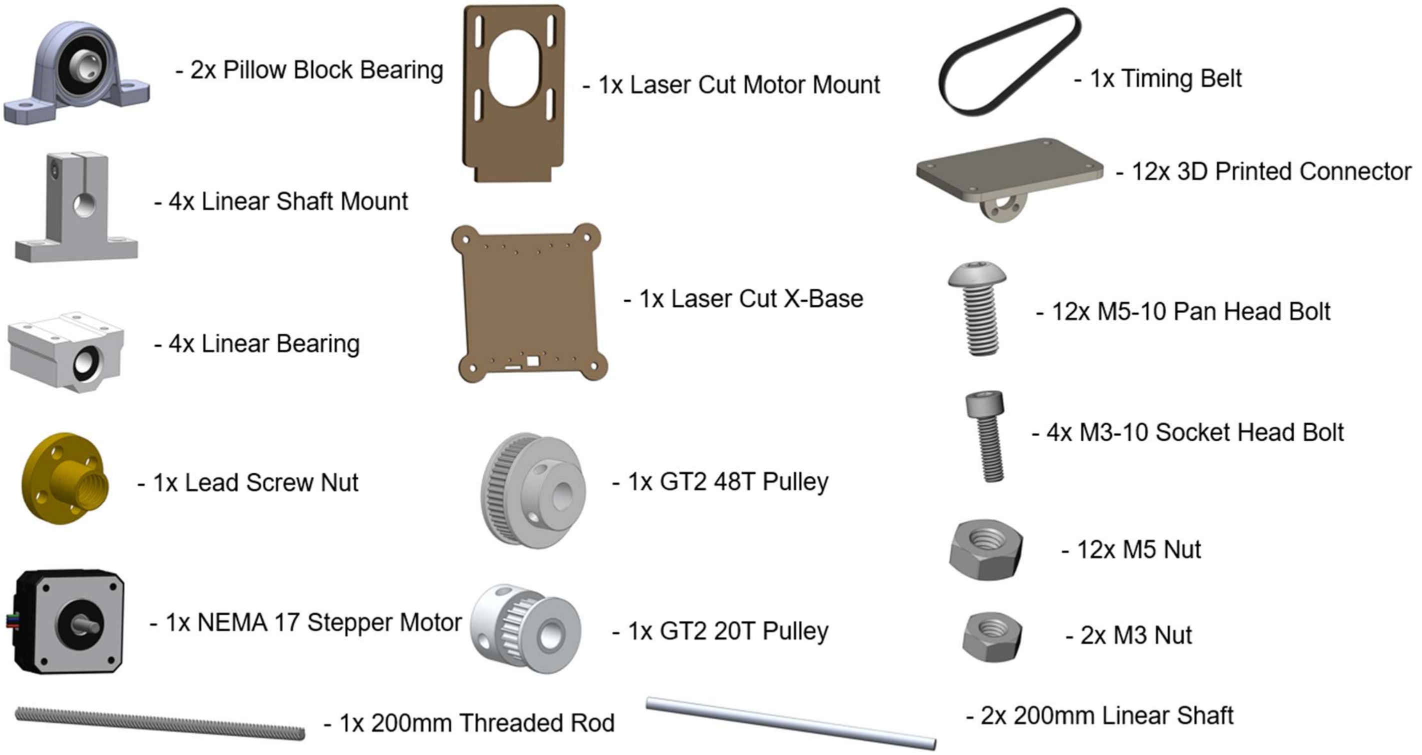


| | | |
|---|---|---|
| 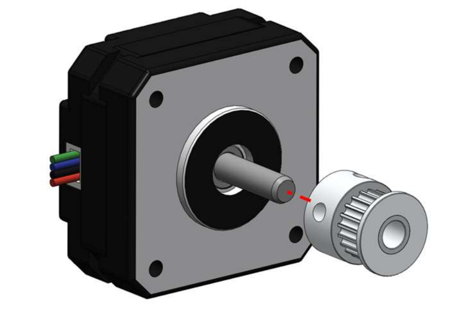 | 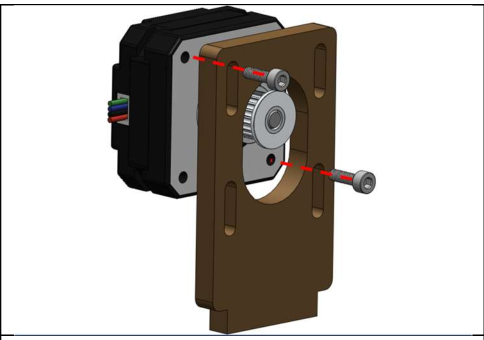 | 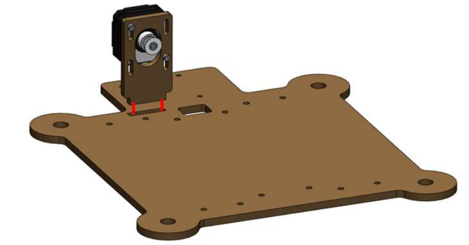 |
| 1. Slide the 20T pulley onto stepper motor shaft. Tighten the two set screws on the pulley. | 2. Screw the stepper motor onto the laser cut motor mount. | 3. Glue the motor mount into the laser cut x-axis base. Use super glue or epoxy. Let fully cure. |
| 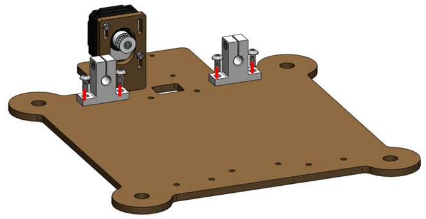 | 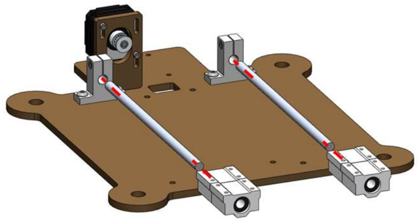 | 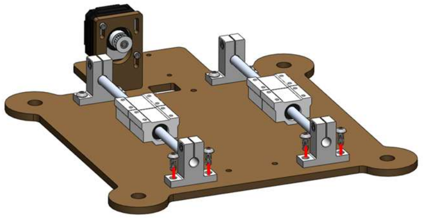 |
| 4. Screw two linear shaft mount onto the base with M5 bolts. | 5. Slide the linear shafts into the mounts and slide two bearings onto each shaft. | 6. Screw two linear shaft mount onto the base with M5 bolts. Tighten the set screws in all linear shaft mounts. |

| | | |
|---|---|---|
| 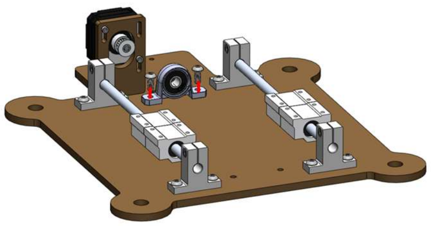 | 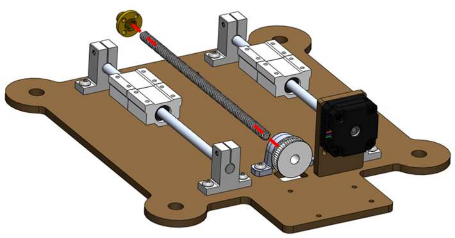 | 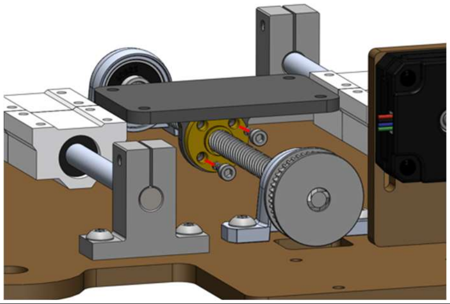 |
| 7. Screw a pillow block bearing onto the base with M5 bolts. | 8. Place the belt on the 42T pulley and slide the threaded rod through the pillow block bearing and the pulley. Spin the lead screw nut onto the rod. | 9. Slide the XY connector onto the threaded rod and secure onto the lead screw nut with 2 M3 bolts and 2 M3 nuts. |
| 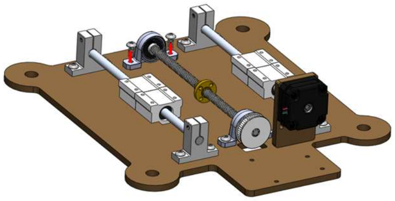 | 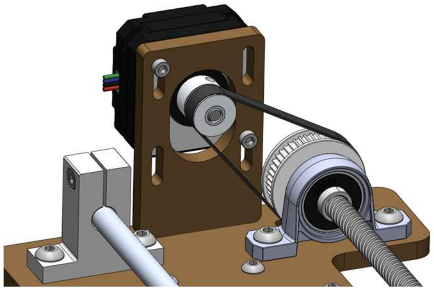 | |
| 10. Screw a pillow block bearing onto the base with M5 bolts. Tighten the set screws on the pillow block bearings. | 11. Adjust the motor mount screws to maintain slight tension in the belt. | |

#### *E. Overall Assembly*

##### Parts

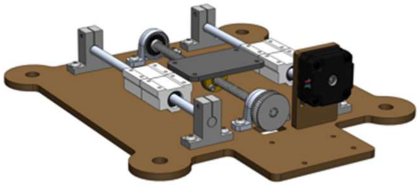

- 1x X-Axis Module

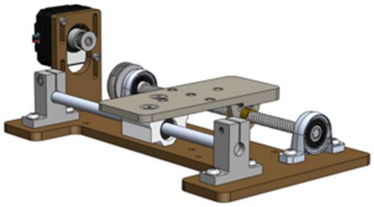

- 1x Y-Axis Module

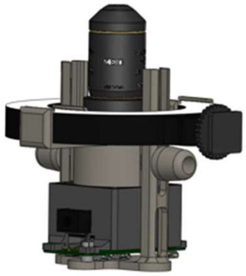

- 1x Z-Axis Stand

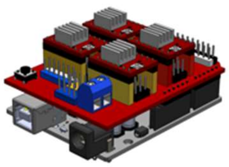

- 1x Arduino Uno + CNC Shield

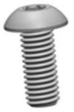

- 14x M4-12 Socket Head Bolt

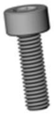

- 4x M2-14 Socket Head Bolt

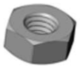

- 4x M3 Nut

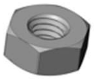

- 4x M4 Nut

| | | |
|---|---|---|
| 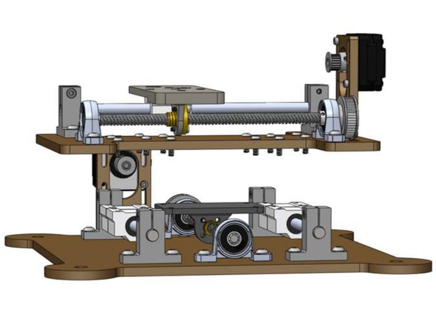 | 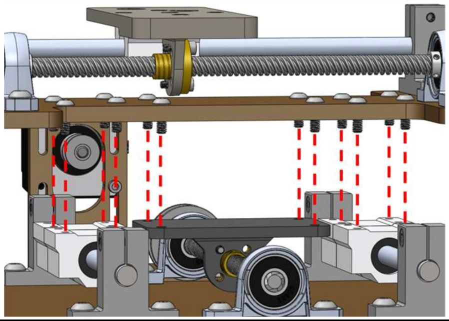 | 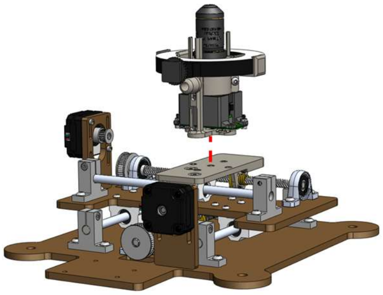 |
| 1. Place 12 M4 bolts into the holes on the y-base. If needed place the included spacers between the x and y axes. | 2. Screw the bolts into the x-axis linear bearings. Back the bolts in the xy-connector with M4 bolts. | 3. Snap the optics z-axis stand onto the y-axis. |
| 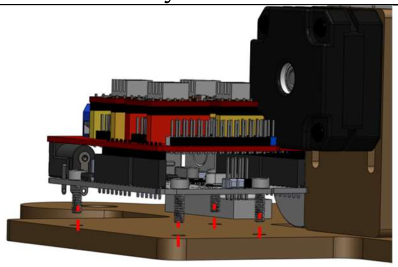 | 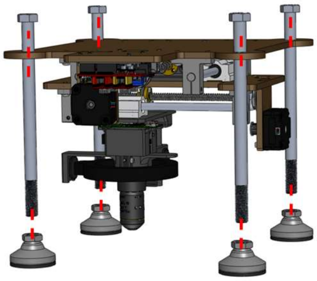 | |
| 4. Screw the Arduino Uno to the x-base with 4 M3 bolts. | 5. Add support legs to the actuator if needed. | |
